# Chikungunya virus vaccine candidate incorporating synergistic mutations is attenuated and protects against virulent virus challenge

**DOI:** 10.1101/2021.10.19.464869

**Authors:** Anthony J. Lentscher, Nicole McAllister, Kira A. Griswold, James L. Martin, Olivia L. Welsh, Danica M. Sutherland, Laurie A. Silva, Terence S. Dermody

## Abstract

**Background:** Chikungunya virus (CHIKV) is an arbovirus that periodically reemerges to cause large epidemics of arthritic disease. While the robust immunity elicited by live-attenuated virus (LAV) vaccine candidates makes them attractive, CHIKV vaccine development has been hampered by a high threshold for acceptable adverse events.

**Methods:** We evaluated the vaccine potential of a recently described LAV, SKE, that exhibits diminished replication in skeletal muscle cells due to insertion of target sequences for skeletal muscle-specific miR-206. We also evaluated whether these target sequences could augment safety of a LAV encoding a previously described attenuating mutation, E2 G82R, which on its own was too reactogenic in clinical trials. Attenuation of viruses containing these mutations was compared with a double mutant, SKE G82R.

**Results:** SKE was attenuated in both immunodeficient and immunocompetent mice and induced a robust neutralizing antibody response, indicating its vaccine potential. However, only SKE G82R elicited diminished swelling in immunocompetent mice at early time points post-inoculation, indicating that these mutations synergistically enhance safety of the vaccine candidate.

**Conclusions:** These data suggest that restriction of LAV replication in skeletal muscle enhances tolerability of reactogenic vaccine candidates and may improve the rational design of CHIKV vaccines.

## INTRODUCTION

Vaccines are a potent defense against viruses with pandemic potential. Central to the development of new vaccines is the balance between safety and immunogenicity. Effective vaccines should provide sufficient antigen at appropriate sites in the host to elicit a protective response and simultaneously avoid vaccine-associated adverse events. Inactivated and subunit vaccines frequently meet the latter criterion, as they are incapable of infection and replication characteristic of live, attenuated vaccines [1]. However, because inactivated and subunit vaccines cannot replicate and disseminate from sites of inoculation, they often require use of adjuvants or multiple doses to promote long-lasting protection [1]. Live-attenuated virus vaccines often elicit protective immune responses following a single dose, but they are more frequently associated with adverse events [1]. It has been especially difficult to achieve the immunogenicity-safety balance for vaccines against chikungunya virus (CHIKV).

CHIKV is a mosquito-transmitted arthritogenic alphavirus that has reemerged to produce devastating epidemics of fever, rash, polyarthralgia, and polyarthritis [2]. CHIKV causes severe disease in as many as 95% of infected individuals [3], and ∼ 60% of those infected will experience chronic joint symptoms lasting months-to-years after initial infection [4]. The virus spreads rapidly following introduction into naïve populations and, although CHIKV infection is rarely fatal in the absence of underlying comorbidities [2], outbreaks pose a significant economic burden both in healthcare costs and loss of work for affected individuals [5]. Exacerbating the threat posed by CHIKV is the absence of approved vaccines and antivirals to limit the global burden of disease.

While the musculoskeletal manifestations of CHIKV disease are severe and often debilitating, the self-limiting nature of virus infection and low rates of CHIKV-associated fatality have raised the threshold for acceptable vaccine-associated adverse events. This high threshold has complicated vaccine development and is best demonstrated in clinical trials of CHIKV strain 181/25, the first live-attenuated CHIKV vaccine candidate to be tested in humans. The vaccine strain was recovered following plaque-to-plaque passage of a clinical isolate until attenuation was empirically achieved based on a small-plaque phenotype in cultured cells and attenuation in newborn mice [6]. In phase II clinical trials, 57 of 58 (∼98%) 181/25-vaccinated individuals developed neutralizing antibodies following a single dose. However, 5 of 59 (∼8%) vaccinees experienced mild, transient arthralgias [7] and, due to this reactogenicity, 181/25 development was halted.

CHIKV strain 181/25 is attenuated by two coding mutations in the viral E2 attachment protein, with the majority of attenuation attributed to a glycine-to-arginine mutation at residue 82 [8–10]. The G82R mutation enhances virus binding to glycosaminoglycans, which are cell-surface carbohydrates that mediate initial attachment of the virus to cells [9,10]. Strain 181/25 elicits a more robust neutralizing antibody response relative to wild-type virus, a phenotype that also segregates with the G82R mutation, and, therefore, is more readily cleared following infection [11]. Because attenuation of 181/25 is attributable to only two coding mutations, reactogenicity in clinical trials is likely due to reversion [12]. Limiting the reversion potential of 181/25 by incorporation of additional attenuating mutations may promote safety without compromising the potent immunogenicity displayed by this vaccine candidate.

We engineered a CHIKV variant, SKE, that is attenuated due to diminished replication in skeletal muscle cells by incorporation of target sequences for skeletal muscle-specific miR-206. SKE infection results in diminished swelling in the inoculated footpad of immunocompetent mice compared with virulent parental strain SL15649 [13]. While both SKE and SL15649 replicate with similar kinetics and reach comparable peak titers in musculoskeletal tissues of mice, SKE infection is associated with reduced infiltration of T cells into skeletal muscle and diminished production of IL-6, a critical mediator of CHIKV-induced disease [13]. We hypothesized that muscle-specific miR-206 target sequences placed in the context of other attenuating mutations will improve safety and retain immunogenicity of live-attenuated CHIKV vaccine candidates.

Here, we investigated whether muscle-specific miR-206 target sequences introduced into a virus containing the G82R mutation of 181/25 could improve safety of a vaccine strain without compromising its immunogenicity. We assessed the virulence and protective capacity of CHIKV strains containing E2 G82R, miR-206 target sequences, or both. We found that vaccination of immunocompetent mice with any of the vaccine candidate strains elicited a robust and protective immune response. However, only mice infected with a virus encoding the combination of mutations were protected from early disease manifestations. Additionally, in immunodeficient mice, only those inoculated with the combination vaccine candidate survived through the duration of the experiment. These data suggest that incorporation of multiple attenuating mutations enhances the safety of vaccine candidates without compromising immunogenicity and will inform the rational design of vaccine candidates capable of limiting the global health threat posed by CHIKV.

## METHODS

### Cells and Viruses

Baby hamster kidney cells (BHK-21; ATCC CCL-10) were maintained in alpha minimal essential medium (αMEM; Gibco) supplemented to contain 10% fetal bovine serum (FBS, Atlanta Biologicals) and 10% tryptose phosphate. Vero81 cells (ATCC CCL-81) were maintained in αMEM supplemented to contain 5% FBS. All cell maintenance media were supplemented to contain 2 mM L-glutamine (Gibco).

The WT CHIKV strain SL15649 (pMH56) was provided by Mark Heise (University of North Carolina at Chapel Hill). The skeletal muscle-restricted (SKE) strain was engineered as described [13]. G82R and SKE G82R were recovered following site-directed mutagenesis of the SL15649 and SKE infectious clones, respectively, using KOD polymerase (EMD Millipore). Reactions were conducted following the manufacturer’s instructions. Full-length sequences of infectious clone plasmids were confirmed by consensus sequencing. Virus stocks were recovered following linearization of infectious clone plasmids using NotI-HF (NEB), *in vitro* transcription, and electroporation of transcribed RNA into BHK-21 cells as described [13]. All experiments using SL15649 and variant clones were conducted using biosafety level 3 conditions.

### Assessment of CHIKV Replication Kinetics and Viral Plaque Assays

Vero81 cells were adsorbed with CHIKV strains diluted in virus dilution buffer (VDB, RPMI medium with 25 mM HEPES [Gibco] supplemented to contain 1% FBS) at a multiplicity of infection of 0.01 plaque-forming units (PFU)/cell at 37°C for 1 h. The viral inoculum was removed, cells were washed twice with PBS, and complete medium was added. Following incubation at 37°C for various intervals, 10% of the culture supernatant was collected and replaced with fresh medium. Serial 10-fold dilutions of culture supernatants were prepared in VDB and adsorbed to Vero81 cells at 37°C for 1 h. Monolayers were overlaid with agarose, and plaques were visualized following neutral red staining as described [13]. Plaques were enumerated in duplicate and averaged to calculate PFU/ml.

### FFU Assays

Serial 10-fold dilutions of samples in VDB were adsorbed to Vero81 cells at 37°C for 2 h. Monolayers were overlaid with methylcellulose, fixed with methanol, washed, and permeabilized as described [13]. Cells were incubated with CHIKV-specific polyclonal antiserum (ATCC, VR-1241AF) diluted 1:1000 in perm/wash buffer (PBS supplemented to contain 0.1% saponin and 0.1% BSA) at RT for 2 h, washed three times with perm/wash buffer, incubated with a horseradish peroxidase-conjugated goat anti-mouse IgG secondary antibody (SouthernBiotech, 1030-05, 1:2000 dilution) at RT for 1 h, and washed three times with perm/wash buffer. Infected cell foci were visualized following incubation with TrueBlue Substrate (VWR, 95059-168) and enumerated using a CTL Biospot Analyzer as described [13].

### Transfection of miRNA-Mimic siRNAs

Vero81 cells were transfected with 10 nM of siRNAs using Lipofectamine RNAiMAX (Invitrogen) diluted in serum-free OPTI-MEM according to the manufacturer’s instructions. At 16 h post-transfection, cells were adsorbed with CHIKV strains diluted in VDB at an MOI of 5 PFU/cell at 37°C for 1 h. The virus inoculum was removed, cells were washed twice with PBS, complete medium was added, and cells were incubated at 37°C for 6 h. Cells were fixed and stained by indirect immunofluorescence to detect CHIKV antigen and cell nuclei as described [14]. Four fields of view per well were imaged using a LionHeart FX automated microscope (BioTek). The percentage infectivity was determined by dividing the number of virus-infected cells by the total number of cells per field of view.

### Mouse Experiments

C57BL/6J mice were obtained from the Jackson Laboratory. *Ifnar*^*−/−*^ mice were provided by John Williams (University of Pittsburgh). Three-to-four-week-old mice were used for all studies. Mice were inoculated in the left rear footpad with 10 μL of diluent alone (PBS supplemented to contain 1% FBS; mock) or diluent containing 10^3^ PFU of each virus. C57BL/6 mice were weighed at 24-hour intervals and monitored for signs of disease. The area of the left rear footpad was determined prior to inoculation by measurement of footpad width and thickness using digital calipers and at 24-hour intervals thereafter for 10 days. At experimental endpoints, mice were euthanized by exposure to isoflurane followed by thoracotomy. Blood was collected, mice were perfused by intracardiac inoculation of PBS, and tissues were resected and homogenized using a MagNA Lyser (Roche) in PBS supplemented to contain 1% FBS prior to determination of viral titer by FFU assay. *Ifnar*^*−/−*^ mice were monitored twice daily for 10 days post-inoculation and euthanized by exposure to isoflurane followed by thoracotomy when weight loss exceeded 15% or at the study endpoint.

All mouse infection studies were conducted using animal biosafety 3 (ABSL3) facilities and guidelines. All animal husbandry and experimental procedures were performed in accordance with U.S. Public Health Service policy and approved by the University of Pittsburgh IACUC.

### Immunization Protocol

C57BL/6J mice were inoculated in the left rear footpad with 10 μl diluent alone (mock) or diluent containing 10^3^ PFU of SL15649, SL15649 G82R, SKE, or SKE G82R. Mice were monitored for weight loss and allowed to recover for 30 d, at which time all mice were challenged by inoculation in the left rear footpad with 10^3^ PFU of SL15649. Mice were monitored for weight loss and footpad swelling for 14 d following challenge.

### Focus-Reduction Neutralization Tests

C57BL/6J mice were inoculated in the left rear footpad with 10 μL diluent alone (mock) or diluent containing 10^3^ PFU of SL15649, SL15649 G82R, SKE, or SKE G82R. At day 30 post-inoculation, mice were euthanized and serum was collected. Serum was heat inactivated at 56°C for 30 min and serially diluted in VDB at a ratio of 1:4. Dilutions were incubated with 100 FFU of SL15649 at 37°C for 1 h. Vero81 cell monolayers were inoculated with antibody-virus mixtures and incubated for 18 h, after which time virus-infected foci were enumerated using the FFU protocol described above. Serum-free controls were used to determine percent neutralization, and the concentration at which 50% of SL15649 was neutralized by immune serum (FRNT_50_) was determined by non-linear regression. Mice that did not exhibit any neutralizing antibody titer at day 30 post-inoculation were excluded from analysis.

### Statistical Analysis

All statistical tests were conducted using GraphPad Prism 9 software. Significant differences were detected using log-rank test or ANOVA with Tukey’s post hoc test to correct for multiple comparisons. *P* values of less than 0.05 were considered to be statistically significant. Descriptions of the specific statistical tests used for each experiment are provided in figure legends.

## RESULTS

### Recovery of CHIKV Vaccine Candidates

CHIKV mutants in this study were engineered into the background of virulent strain SL15649. To determine whether combination of the E2 G82R mutation with target sequences for skeletal muscle-specific miR-206 present in SKE enhances the safety of a vaccine candidate without diminishing protective efficacy, we recovered strain SKE G82R using site-directed mutagenesis (Figure 1). To control for contributions of the E2 G82R mutation alone, we also recovered SL15649 containing the G82R mutation (SL15649 G82R). SKE was recovered as described previously and used to define attenuation mediated by miR-206 restriction [13]. Hereafter, SL15649 G82R, SKE, and SKE G82R will be referred to collectively as vaccine candidate strains. All experiments were conducted alongside wild-type SL15649 as a virulent virus control.

**Figure 1.**
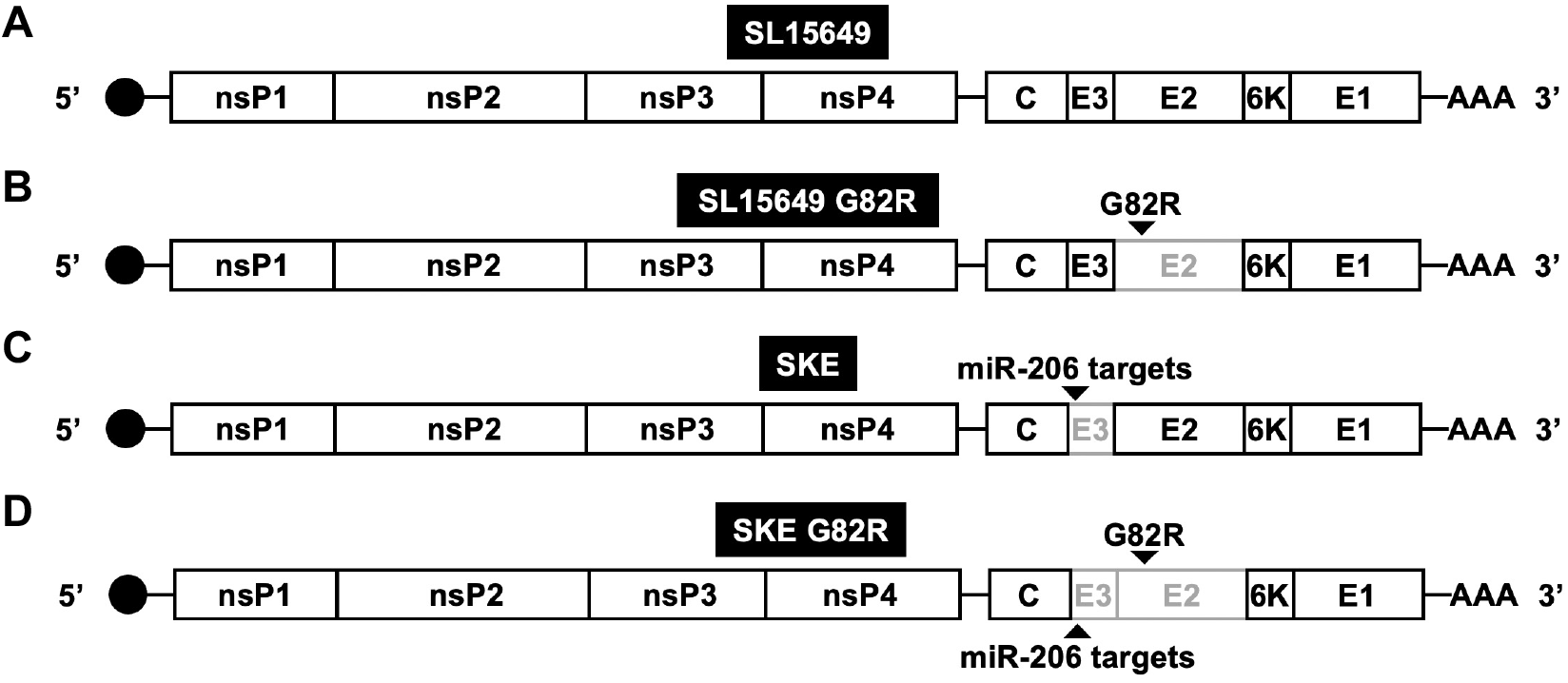
Schematic of vaccine candidate CHIKV strains. Vaccine strains were engineered into the background of virulent CHIKV strain SL15649 (*A*). Vaccine strains were recovered containing a G82R mutation in the viral E2 protein (SL15649 G82R; *B*), incorporating target sequences for skeletal muscle-specific miR-206-3p in the E3 coding region (SKE; *C*), or both (SKE G82R; *D*).

### Characterization of Vaccine Candidate Strains Using Cultured Cells

We first assessed the capacity of CHIKV vaccine candidate strains to infect Vero81 cells, which do not naturally express miR-206 (data not shown). Cells were adsorbed at an MOI of 0.01 PFU/cell, and supernatants were harvested at various intervals post-adsorption to quantify viral progeny production by plaque assay. All viruses replicated to high titer in these cells (Figure 2A).

**Figure 2.**
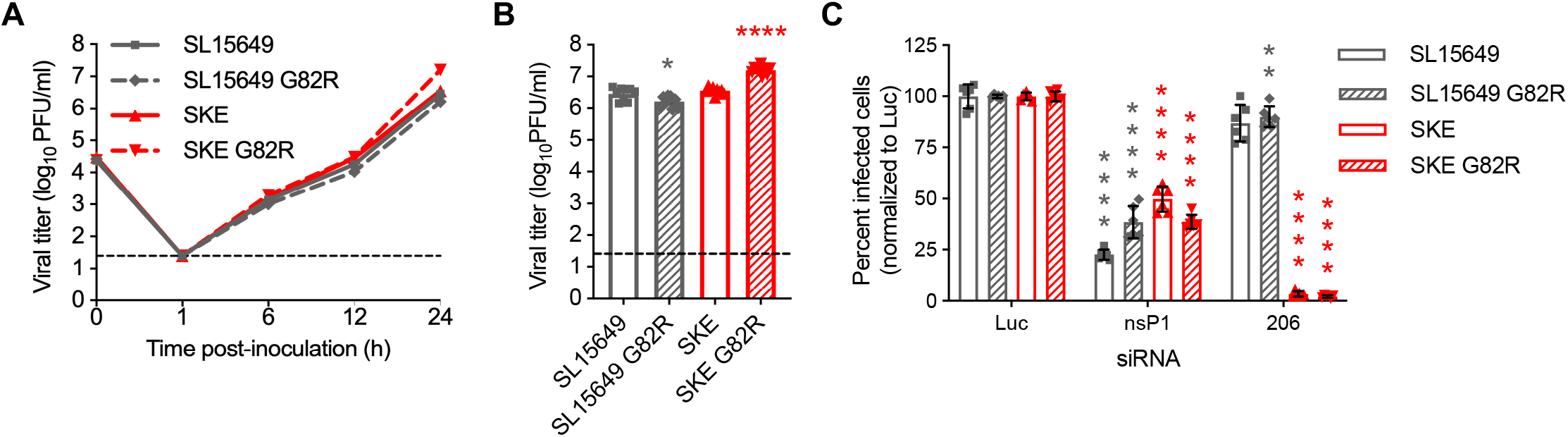
Replication of vaccine candidate CHIKV strains in cultured cells. *A*, Vero81 cells were adsorbed with SL15649, SL15649 G82R, SKE, or SKE G82R at an MOI of 0.01 PFU/cell. Supernatants were collected at the times shown post-adsorption, and viral titer was quantified by plaque assay. Data gathered at 24 h post-inoculation (*B*) are displayed separately to show significant differences more clearly. Results are expressed as the mean viral titer from duplicate wells for three independent experiments. Error bars indicate SEM. Dashed lines indicate the limit of detection. *P* values were determined comparing vaccine candidate strain replication to SL15649 at 24 h post-inoculation by ANOVA followed by Tukey’s post hoc test. * *P* < 0.05; **** *P* < 0.0001. *C*, Vero81 cells were transfected with siRNA directed against luciferase or CHIKV nsP1 or muscle-specific miR-206-mimic siRNA and adsorbed with SL15649, SL15649 G82R, SKE, or SKE G82R at an MOI of 5 PFU/cell. Cells were fixed at 6 h post-adsorption, and the percentage of CHIKV-infected cells was determined by indirect immunofluorescence. Results are expressed as the mean percentage of infected cells normalized to luciferase control siRNA from three triplicate wells for two independent experiments. Error bars indicate SEM. *P* values were determined comparing nsp1 and mir-206 siRNA groups to luciferase controls for each virus by ANOVA followed by Tukey’s post-hoc test. ** *P* < 0.01; **** *P* < 0.0001.

To determine susceptibility of vaccine candidate strains to miR-206-mediated restriction, we transfected Vero81 cells with an siRNA targeting the viral nsP1 gene or an siRNA mimicking the sequence of miR-206. Transfected cells were adsorbed with WT virus or vaccine strains at an MOI of 5 PFU/cell and fixed at 6 h post-adsorption. The percentage of infected cells was quantified by indirect immunofluorescence. While all viruses were restricted by the nsP1 siRNA, only SKE and SKE G82R were potently restricted by the miR-206-mimic siRNA (Figure 2C). These data demonstrate that vaccine candidate strains recovered for this study replicate in cultured cells with anticipated characteristics.

### SKE G82R Displays Greater Attenuation in Mice than SKE and SL15649 G82R

To define safety of the CHIKV vaccine candidate strains, we used an acute model of CHIKV disease in which C57BL/6J mice are inoculated in the left rear footpad with 1000 PFU of virus, and swelling of the infected foot is quantified using digital calipers. Mice inoculated with SL15649 experienced biphasic swelling, peaking at days 3 and 7 post-inoculation (Figure 3A), which is characteristic of CHIKV disease in C57BL/6J mice [13,15]. The vaccine candidate strains elicited less swelling at day 7 post-inoculation, demonstrating attenuation of these viruses in immunocompetent mice (Figure 3C). Although swelling in mice inoculated with all vaccine strains trended lower at day 3 post-inoculation than in mice inoculated with WT virus, only those inoculated with SKE G82R displayed significantly less swelling, indicating that mutations incorporated into SKE G82R synergistically diminish virulence in these animals (Figure 3B).

**Figure 3.**
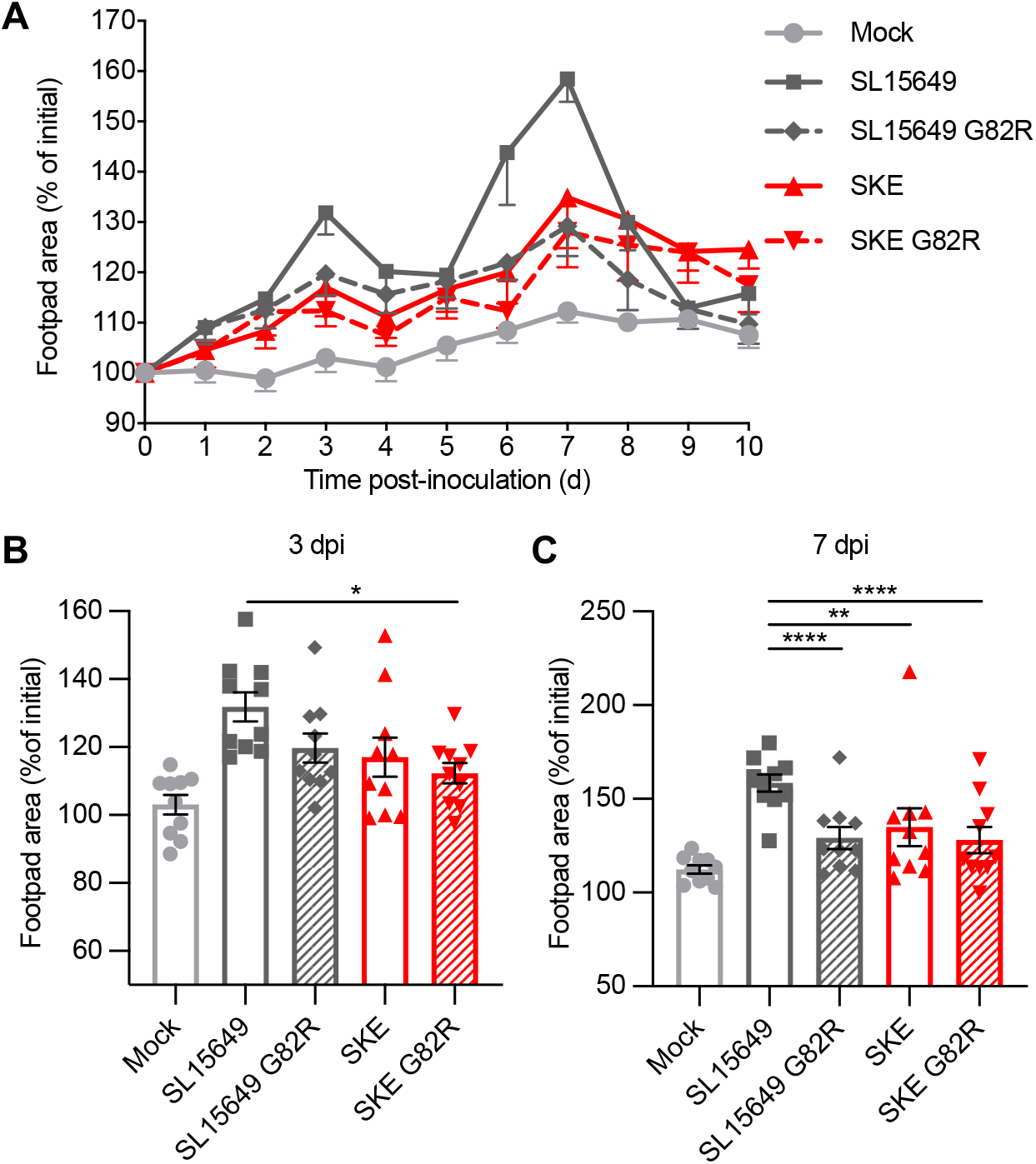
CHIKV vaccine candidate strains are attenuated in immunocompetent mice. *A*, Three-to-four-week-old C57BL/6J mice were inoculated in the left rear footpad with 10^3^ PFU of SL15649, SL15649 G82R, SKE, or SKE G82R. Left rear footpad swelling was quantified using digital calipers on the days shown post-inoculation. Data at day 3 (*B*) and 7 (*C*) post-inoculation are displayed separately to show significant differences more clearly. *A-C*, Results are normalized to initial footpad area and presented as the mean percentage of initial footpad area for 10 mice per group. Error bars indicate SEM. *P* values were determined by ANOVA followed by Tukey’s post hoc test. * *P* < 0.05; ** *P* < 0.01; **** *P* < 0.0001.

### Vaccine Candidate Strains Are Attenuated in Immunocompromised Mice

Because safety is a principal concern for live-attenuated vaccine candidates, we assessed virulence of the CHIKV vaccine candidate strains in experiments using immunocompromised mice. Mice lacking expression of the interferon-α/β receptor (IFNAR) were inoculated with 1000 PFU of virus and monitored for survival for 10 d post-inoculation. Relative to infection with SL15649, all vaccine candidate strains were attenuated following infection of IFNAR-deficient mice (Figure 4). However, the only virus-infected mice that survived until the end of the experiment were infected with SKE G82R, suggesting attenuation of that strain in immunodeficient animals.

**Figure 4.**
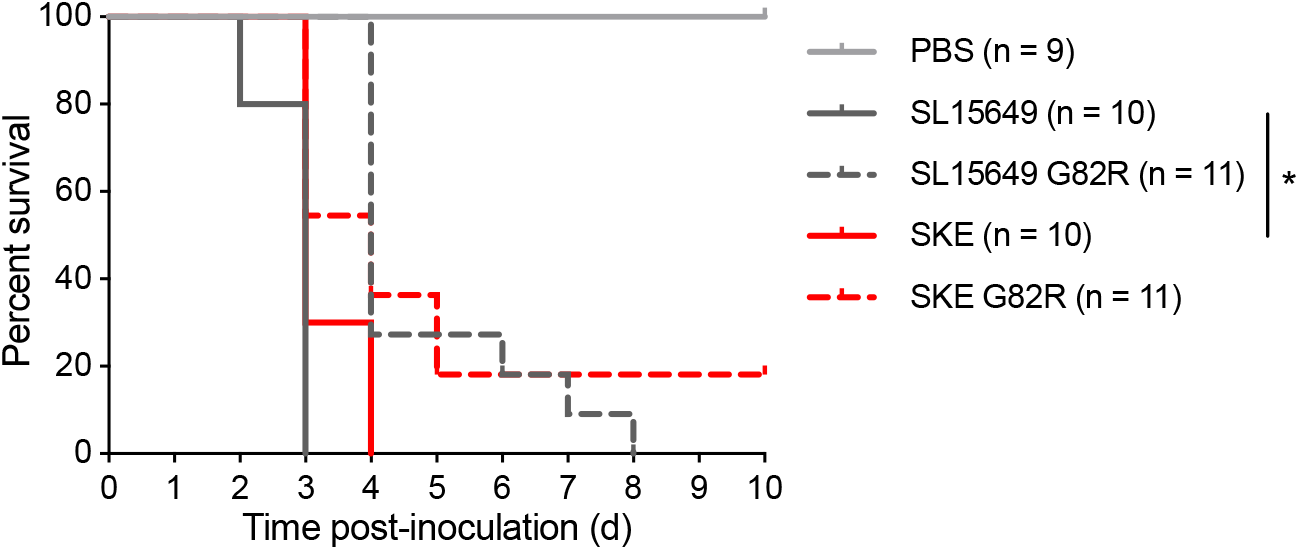
CHIKV vaccine candidate strains are attenuated in immunocompromised mice. Four-week-old *Ifnar*^*−/−*^ mice were inoculated in the left rear footpad with 10^3^ PFU of SL15649, SL15649 G82R, SKE, or SKE G82R. Survival was assessed for 10 days post-inoculation (n = 9 to 11 mice per group). *P* values were determined by log-rank test comparing SL15649 with SKE and SL15649 G82R with SKE G82R. * *P* < 0.05.

### Immune Responses Elicited by Vaccine Candidate Strains Protect Immunocompetent Mice from Disease Following Virulent Virus Challenge

To define the protective capacity of the CHIKV vaccine candidate strains, we inoculated mice with PBS (mock) or 1000 PFU of SL15649 or each vaccine candidate strain and challenged the animals 30 days later by inoculating 1000 PFU of SL15649. Footpad swelling was quantified in the inoculated footpad using digital calipers. While mock-vaccinated mice displayed biphasic swelling, none of the mice inoculated initially with SL15649 or the vaccine candidate strains developed swelling in the inoculated foot (Figure 5). Thus, each of the CHIKV vaccine candidate strains protects mice against virulent virus challenge.

**Figure 5.**
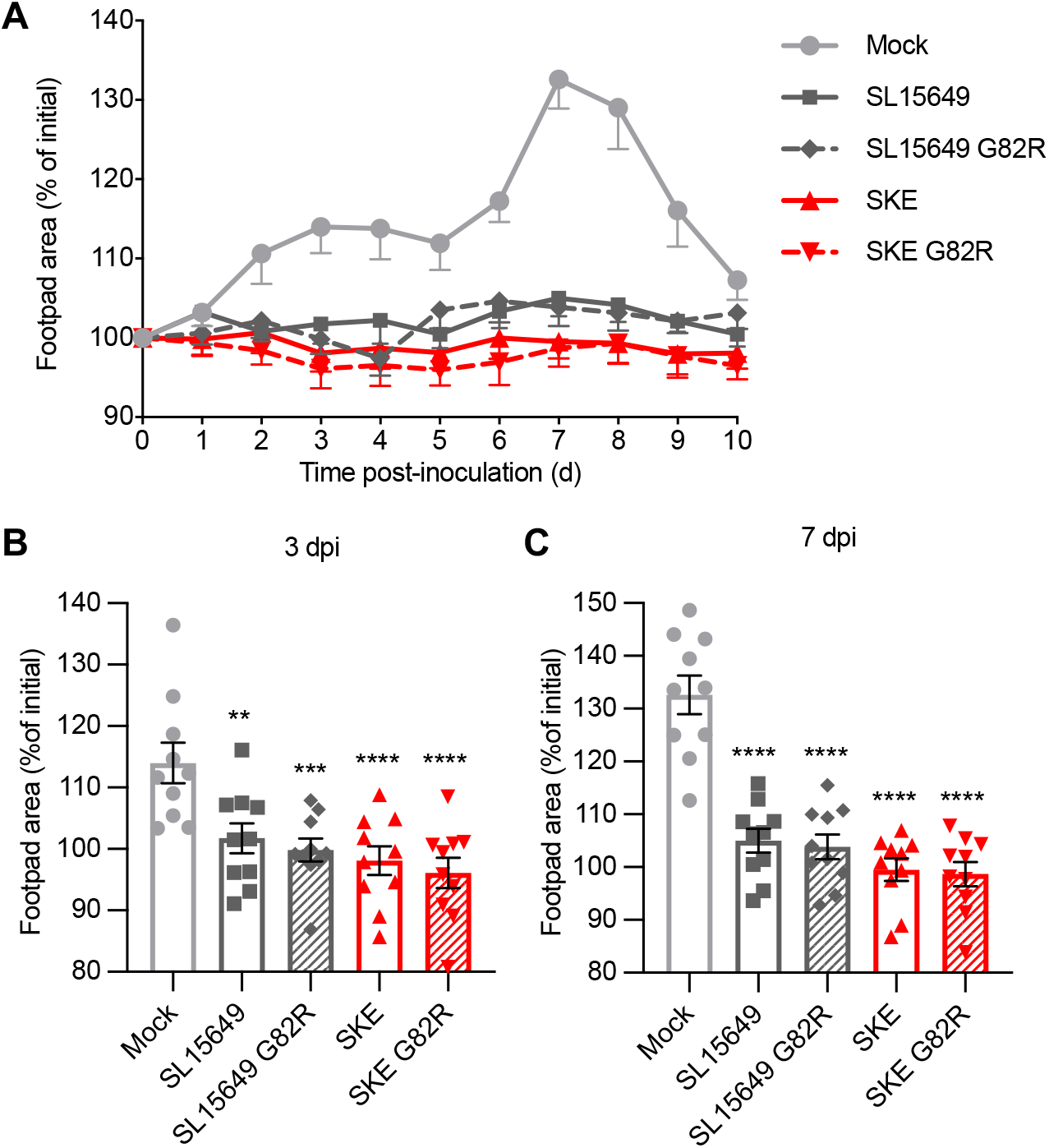
Inoculation of immunocompetent mice with CHIKV vaccine candidate strains leads to protection against virulent virus challenge. Three-to-four-week-old C57BL/6J mice were inoculated in the left rear footpad with 10^3^ PFU of SL15649, SL15649 G82R, SKE, or SKE G82R. At day 30 post-inoculation, mice were challenged in the left rear footpad with 10^3^ PFU of SL15649. Left rear footpad swelling was quantified using digital calipers on the days shown post-challenge. Data at day 3 (*B*) and 7 (*C*) post-challenge are displayed separately to show significant differences more clearly. *A-D*, Results are normalized to initial footpad area and presented as the mean percentage of initial footpad area for 10 mice per group. Error bars indicate SEM. *P* values were determined by ANOVA followed by Tukey’s post hoc test. Statistically significant differences were found in comparing vaccinated groups to mock-vaccinated mice. ** *P* < 0.01; *** *P* < 0.001; **** *P* < 0.0001.

### SL15649 G82R Titers Are Diminished in the Left Ankle at 3 Days Post-Inoculation

Live-attenuated vaccines often elicit robust, durable immune responses due to the capacity of the attenuated virus to replicate and disseminate to sites targeted by the virulent counterpart. To determine whether the CHIKV vaccine candidate strains replicate at the site of inoculation and disseminate to sites of secondary replication, we inoculated C57BL/6J mice in the left rear footpad with 1000 PFU of virus and resected various tissues at day 3 post-inoculation. CHIKV titers in the left and right ankles were quantified by focus-forming unit (FFU) assay, and titers in the left quadricep muscle were quantified by RT-qPCR. While SKE and SKE G82R reached similar titers in the left ankle, which is adjacent to the inoculation site, titers of SL15649 G82R were diminished at this time point (Figure 6A). Virus was undetectable in the left ankle in two of five SL15649 G82R-infected mice, indicating that this strain either replicates inefficiently or is rapidly cleared from this site. The mice in these experiments received comparable inocula, as judged by detection of virus in the left quadricep muscle of all virus-infected mice. Additionally, although titers in the right ankle of SL15649 G82R-infected mice did not differ significantly from those in mice inoculated with the other vaccine candidate strains, virus was undetectable in four of five SL15649 G82R-infected mice, indicating that this strain does not disseminate efficiently to musculoskeletal sites of secondary replication by day 3 post-inoculation.

**Figure 6.**
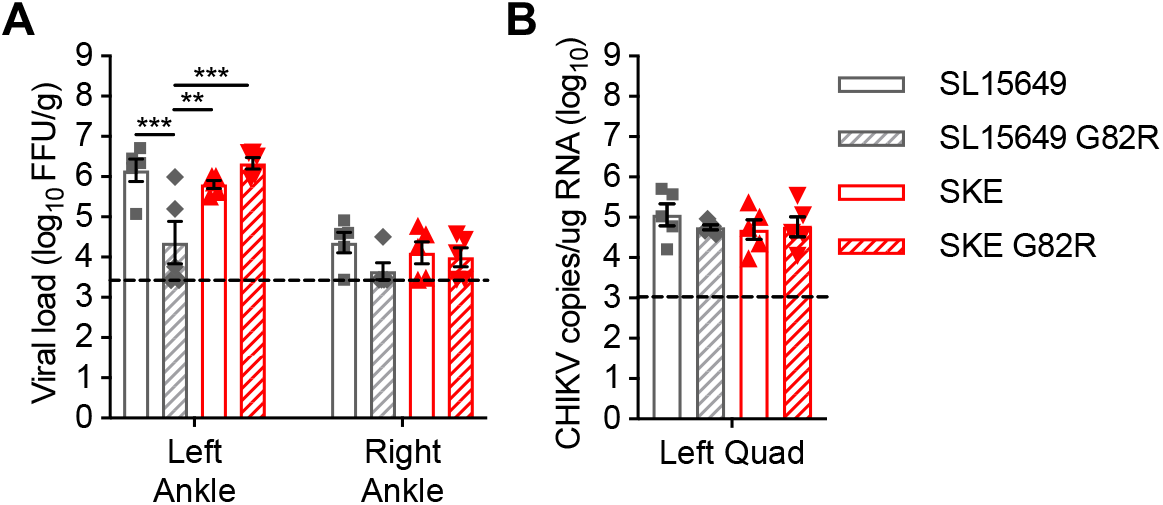
Titers of SL15649 G82R are diminished in the left ankle of immunocompetent mice. Three-to-four-week-old C57BL/6J mice were inoculated in the left rear footpad with 10^3^ PFU of SL15649, SL15649 G82R, SKE, or SKE G82R. At day 3 post-inoculation, mice were euthanized, and ankles and quadricep muscles were excised. Viral titers in ankle homogenates were determined by FFU (*A*) and in left quadriceps muscle tissue by RT-qPCR (*B*). Horizontal bars indicate mean FFU/g (*A*) or mean CHIKV RNA copies/mg RNA (*B*) for 5 mice per group. Error bars indicate SEM. *P* values were determined by ANOVA followed by Tukey’s post hoc test. ** *P* < 0.01; *** *P* < 0.001.

### Vaccine Candidate Strains Elicit Neutralizing Antibody Responses Comparable to Infection with WT SL15649 in Immunocompetent Mice

Neutralizing antibodies elicited by WT CHIKV are a critical component of a protective antiviral response [16,17]. To determine whether mutations introduced into vaccine candidate strains alter the capacity to elicit a neutralizing antibody response, we inoculated C57BL/6J mice in the left rear footpad with 1000 PFU of virus. At day 30 post-inoculation, serum was collected to quantify neutralizing antibody titers by FFU assay. Infection with vaccine candidate strains elicited neutralizing antibody responses of similar magnitude to SL15648 (Figure 7), indicating that mutations incorporated into the vaccine candidate strains do not diminish immunogenicity.

**Figure 7.**
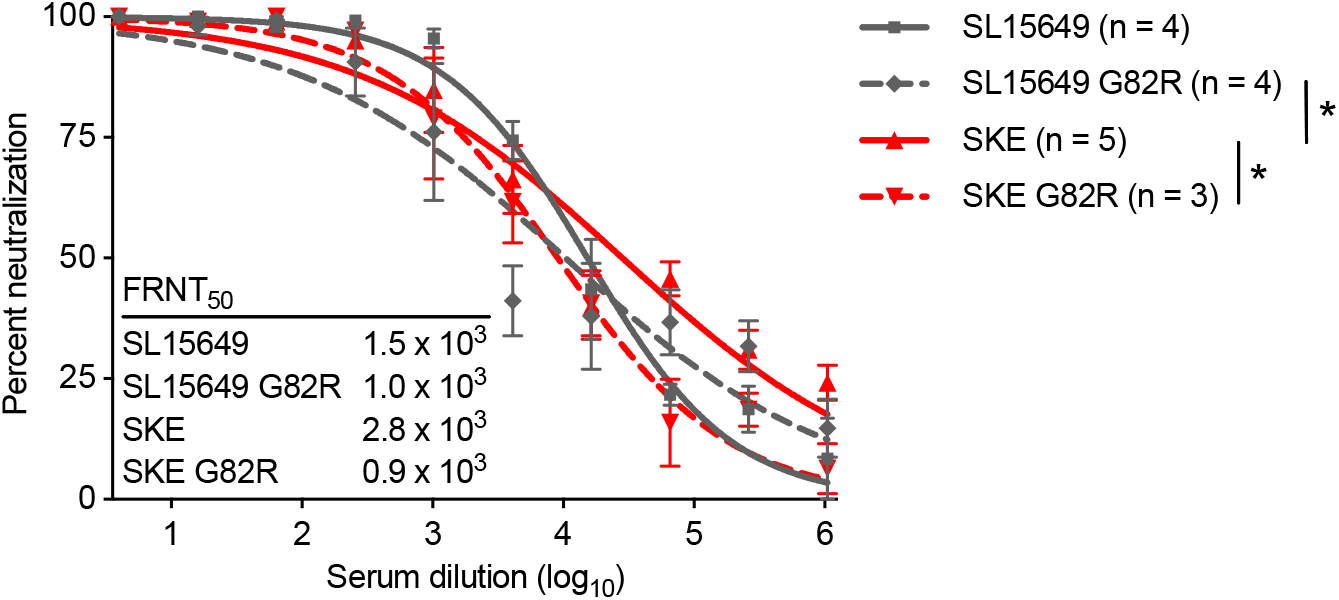
Serum from vaccinated immunocompetent mice efficiently neutralizes CHIKV infection. Three-to-four-week-old C57BL/6J mice were inoculated in the left rear footpad with PBS (mock) or 10^3^ PFU of SL15649, SL G82R, SKE, or SKE G82R. At day 30 post-inoculation, mice were euthanized. Serum was collected and heat inactivated at 56°C for 30 min. Serial dilutions (1:4) of serum were incubated with 100 FFU of SL15649 at 37°C for 1 h. Vero81 cells were inoculated with antibody-virus mixtures and incubated for 18 h, after which time virus-infected foci were enumerated. The concentration at which 50% of SL15649 was neutralized by immune serum was determined using non-linear regression. Mice that did not exhibit any neutralizing antibody titer at day 30 post-inoculation were excluded from analysis. *P* values were determined by ANOVA followed by Tukey’s post hoc test. * *P* < 0.05.

## DISCUSSION

A high threshold for vaccine-associated adverse events has made developing a CHIKV vaccine that is both safe and effective difficult. In this study, we assessed the potential for combining multiple attenuating mutations into a single vaccine strain to optimize this balance. We recovered a virus engineered to contain a previously identified attenuating mutation, E2 G82R, in combination with target sequences for skeletal muscle cell-specific miR-206, as well as control viruses containing each mutation individually. All vaccine candidates elicited diminished swelling in immunocompetent mice relative to virulent CHIKV at day 7 post-inoculation. Interestingly, only the SKE G82R strain, which contains both attenuating mutations, was attenuated at day 3 post-inoculation, indicating that the double mutant displays enhanced safety in mice. All vaccine candidates elicited a neutralizing antibody response in immunocompetent mice and were attenuated in IFNAR-deficient mice. However, only IFNAR-deficient mice infected with SKE G82R survived to day 10 post-inoculation.

While the G82R mutation has been more extensively characterized in the context of vaccine development with 181/25, we found that G82R also is attenuating in the context of strain SL15649. Incorporation of muscle-specific miR-206 target sequences represents a new vaccine strategy for CHIKV, as viruses containing these mutations elicit marked attenuation in mice without compromising tissue titers and antigen availability during acute infection [13]. Although SKE attenuation presumably is due mainly to diminished muscle cell necrosis and recruitment of T cells into infected muscle during the adaptive immune response to infection [13], significant differences in survival of IFNAR-deficient mice infected with SKE versus WT SL15649 indicates an innate mechanism of attenuation. This attenuation could be due to diminished production of proinflammatory mediators [13] or an undefined mechanism. The SL15649 G82R and SKE G82R candidate strains retained virulence in IFNAR-deficient mice, which is unsurprising considering that 40% of IFNAR-deficient mice infected with 181/25 succumb to disease when inoculated with 10 PFU of virus [18]. These data provide evidence that multiple attenuating mutations may be required to yield a vaccine capable of achieving licensure for use in humans.

Other live-attenuated variants of CHIKV have been developed that may comprise a toolkit for production of an optimal vaccine candidate. An interesting candidate, CHIKV-IRES, was recovered following replacement of the subgenomic promoter with an internal ribosomal entry sequence (IRES) [19]. While this virus is safe in immunodeficient mice, it also displays a severe replication defect *in vivo* [19], which may limit immunogenicity in humans. A promising live-attenuated CHIKV vaccine candidate, Δ5nsP3 (also called VLA1553), was recovered following deletion of 183 base pairs of the viral nsP3 gene [20]. This virus elicits a protective neutralizing antibody response in immunocompetent mice [20] and nonhuman primates [21]. In phase I clinical trials, the vaccine was found to be safe and well tolerated in low- and medium-dose groups, but high-dose inoculation was associated with a higher reactogenicity profile [22]. Notable systemic adverse events across all groups included fever, headache, fatigue, and muscle pain [22]. VLA1553 is currently in phase 3 clinical trials. If it is deemed to reactogenic for licensure, further attenuation by restriction of viral replication in muscle cells may improve its safety profile. In mice, CHIKV replication in skeletal muscle promotes release of proinflammatory mediators and muscle damage [13]. If these effects occur following CHIKV infection in humans, limiting muscle replication may diminish the fever and muscle pain observed in phase I trials of VLA1553.

Our study fosters the concept of a vaccine toolbox to inform the rational design of live-attenuated vaccine candidates, especially for viruses like CHIKV, for which a safe and effective vaccine has been elusive. While several live-attenuated CHIKV variants have been described, variants attenuated by different mechanisms to modulate virulence and immunogenicity, such as the SKE G82R vaccine candidate described here, offer an important advantage. Synthesis of knowledge gained from studies of CHIKV pathogenesis may be important to achieving licensure of a vaccine to limit the burden of CHIKV disease.

## NOTES

### Acknowledgments

The authors thank Pavithra Aravamudhan and Gwen Taylor for critically reviewing the manuscript. We are grateful to members of the Silva and Dermody laboratories for useful discussions during the conduct of these studies. We thank John Williams (University of Pittsburgh) for providing Ifnar^−/−^ mice.

### Disclaimer

The findings and conclusions in this report are those of the authors and have not been influenced by the funders.

### Financial support

This work was supported by Public Health Service awards T32 AI049820 (A.J.L.), T32 AI60525 (N.M.), F31 AI147440 (N.M.), and R01 AI123348 (T.S.D.). Additional funding was provided by UPMC Children’s Hospital of Pittsburgh (L.A.S.) and the Heinz Endowments (T.S.D.).

### Author contribution

A.J.L., N.M., L.A.S., and T.S.D. designed research studies. A.J.L., N.M., K.G., J.M., O.W., and D.M.S. conducted experiments. A.J.L. and N.M. acquired data. A.J.L., N.M., K.G., J.M., L.A.S., and T.S.D. analyzed data. A.J.L., N.M., L.A.S., and T.S.D. wrote the manuscript.

### Potential conflicts of interest

The authors declare that there are no conflicts of interest.

## Notes

### Competing Interest Statement

The authors have declared no competing interest.

